# PARiS: Probabilistic Assignment and Repartitioning of isomiR Sequences: A data-driven method for denoising isomiR read count data

**DOI:** 10.64898/2026.05.09.723882

**Authors:** Hannah K. Swan, Andrea M. Baran, Ernesto Aparicio-Puerta, Marc K. Halushka, Seong-Hwan Jun, Matthew N. McCall

## Abstract

MicroRNAs (miRNAs) are non-coding RNAs, approximately 18–24 nucleotides in length, with important gene regulatory functions. In small RNA sequencing (sRNA-seq), observed isoforms of miRNA, called isomiRs, arise from my biological and technical processes. Alterations in isomiR expression has been linked to a wide variety of human diseases, from cancers to neurological diseases. However, it is difficult to distinguish between technical and biological isomiRs. We present PARiS, an algorithm for the Probabilistic Assignment and Repartitioning of isomiR Sequences, that identifies technical error isomiRs in sRNA-seq data and reassigns them to their most likely biological source. We assess the ability of PARiS to identify and remove error isomiR sequences in a realistic simulation study. Additionally, we compare PARiS to alternative approaches, focusing on downstream miRNA-level differential expression analysis in a variety of settings, including a set of simulated datasets, an experimental benchmark dataset, and three colorectal adenocarcinoma cell lines.

## Introduction

MicroRNAs (miRNAs) are a subclass of small, non-coding, single-stranded RNA molecules, approximately 18-24 nucleotides in length, with important biological functions. MiRNAs regulate gene expression (1), control cell differentiation (2), and are even used for intercellular communication (3). The expression of miRNAs must be tightly controlled for healthy cell growth and development (1). Aberrant miRNA expression often results in diseased cells. For example, changes in miRNA expression have been associated with many different types of cancer (6, 7), autoimmune diseases (10), cardiovascular diseases (2), and neurological diseases, like Huntington’s disease (3, 13).

MiRNA expression profiles are typically studied using small RNA sequencing (sRNA-seq), which couples high-throughput sequencing technologies with a size selection step to generate reads of miRNA sequences across different experimental conditions. High-throughput sequencing has largely replaced microarray and qPCR-based quantification due to its ability to identify novel miRNAs and to differentiate between sequences at single nucleotide resolution. The use of sRNA-seq led to the detection of novel miRNA isoforms, called isomiRs. Although these isomiRs were initially thought to be the result of technical errors in sequencing and/or alignment, subsequent studies demonstrated that isomiRs were too abundant in samples to be exclusively produced by sequencing errors, suggesting that isomiRs must be products of biosynthesis pathways in various organisms (4). Since this discovery, researchers have shown that isomiRs are biologically functional molecules (5). Furthermore, studies have linked the expression of not just miRNAs, but specific isomiRs to various diseases including obesity, diabetes, Alzheimer’s disease, and cancers (6, 7).

IsomiRs are classified on the basis of how they differ from a canonical or reference miRNA sequence reported in a miRNA database, such as miRBGeneDB (8) or miRBase (9). A detailed nomenclature to describe isomiRs was developed by the miRNA Transcriptomic Open Project (miRTOP) (10). The addition or removal of nucleotides from the 5’ or 3’ end of the reference sequence produces a 5’ isomiR or 3’ isomiR, respectively. In this manuscript, we refer to isomiRs resulting from such changes as length variant isomiRs. The substitution of nucleotides within the reference sequence produces a polymorphic isomiR. We will refer to polymorphic isomiRs in this manuscript as sequence variant isomiRs. Finally, mixed-type isomiRs are the result of both length and sequence differences (11).

Although studies have shown that some isomiR sequences are biologically produced and functional, isomiR sequences are not exclusively biologically produced. During the sequencing process, nucleotides may be erroneously added, deleted, or substituted from a true sequence (19). These random errors produce unwanted technical isomiRs in the sequencing data. Furthermore, these technical variants are indistinguishable from biologically produced isomiRs. Failing to remove these technical isomiRs from miRNA-seq data prior to analysis can affect the calculation of isomiR expression levels and the estimation of differential expression (12). This suggests a need to denoise the isomiR data prior to analysis.

Despite the importance of isomiRs, the most commonly used approach to analyzing miRNA-seq data is to aggregate isomiR-level count data to the miRNA-level after alignment. The aggregation is done by summing read counts for all sequences mapping to the same miRNA. After aggregation, miRNA differential expression is estimated using methods initially developed for messenger RNA sequencing (mRNA-seq) data, such as edgeR (13) or DESeq2 (14). By aggregating both biological and technical isomiRs, this approach substantially reduces the unwanted noise introduced by technical isomiRs. However, aggregating isomiR sequences to the miRNA-level results in a loss of information and potentially obscures important functional differences. Furthermore, recent studies have shown that using isomiR-level expression data to infer miRNA-level differential expression can improve inference (23, 15).

An alternative approach to handling unwanted technical variation in isomiR-level sequencing data is implemented in miREC, a miRNA error RECtification tool (12). MiREC uses a k-mer lattice approach to identify and remove error sequences from the data. Although the model is equipped to handle both substitution errors, resulting in technical sequence variant isomiRs, and addition/deletion errors, resulting in technical length variant isomiRs, assessment of the method focused on its ability to remove substitution errors (12). Currently, there are no benchmark studies that demonstrate that including miREC as a first step in an analytical pipeline for miRNA differential expression improves downstream analysis. Additionally, miREC requires the selection of hyperparameters that may substantially alter performance.

The need to denoise sequencing reads is not unique to miRNA-seq data. In microbiome research, amplicon sequencing is used to quantify variants of a specific marker gene, often the 16S rRNA gene. The marker gene sequence reads are clustered together into Operational Taxonomic Units (OTUs) as a first step in an analytical pipeline for differential abundance analyses. The aggregation of marker gene sequence reads to the OTU level allows the removal of unwanted technical variation from the data, but precludes analysis of the data at a finer resolution than the OTUs. To resolve this, several methods for denoising microbiome data were developed: Deblur (16), DADA2 (17), and UNOISE2 (18). All of these methods infer a set of true marker gene sequences and associated read counts, although the way in which the methods determine membership to the set of true marker gene sequences differs. In the context of miRNA-seq, a given miRNA would be analogous to a marker gene, and the isomiRs of that miRNA would be analogous to the OTUs. However, none of the methods developed for microbial amplicon sequencing data can be directly applied to miRNA-seq data because they focus solely on sequence variants. In fact, Deblur, UNOISE2, and DADA2 all implement a filtering or trimming step to remove length variants from the data. A novel, data-driven approach for denoising isomiR sequence data that can handle all types of isomiR sequences needs to be developed.

To improve the analysis of miRNA-seq data, we propose PARiS, an algorithm for Probabilistic Assignment and Repartitioning of isomiR Sequences. PARiS models the process of generating error isomiR reads from true isomiR sequences based on the observed data and uses it to define a partition of sequences for each miRNA. For a given miRNA, each element of the partition contains a single inferred true isomiR sequence and the set of all error isomiR sequences produced by technical variation around the true isomiR. Denoised isomiR read counts are produced by summing the reads within each element of a partition. We demonstrate the efficacy of PARiS using a combination of simulations, an experimental synthetic benchmark dataset, and three colorectal adenocarcinoma cell lines.

## Materials and Methods

A miRNA profiling experiment typically consists of two or more experimental conditions, with multiple samples within each experimental condition. Let *j* = 1, …, *J* refer to the unique (biological) samples in a miRNA profiling experiment. Each sample consists of isomiR sequences, *s*, with *y*_*sj*_ ∈ ℤ_*≥*0_ referring to the observed read count of isomiR sequence *s* in sample *j*. Each sequence has been mapped to a miRNA *i, i* = 1, …, *I*, where each *i* ∈ ℤ^+^. We assume that there are one or more isomiR sequences *s* mapping to each miRNA *i*. The mapping of isomiR sequence *s* to miRNA *i* is given by *f*(*s*) = *i*, where the function *f* is determined by the choice of alignment software and reference database. We organize the isomiR sequence read counts *y*_*sj*_ in a *K* × *J*-dimensional matrix, *Y* . Here *J* has the same definition as above and *K* is the total number of unique isomiR sequences in the profiling experiment. We organize the additional information from the alignment of the isomiR sequences to the reference database in another data table. The data table with the alignment information has *K* rows and up to 3 columns. The first column is for each unique isomiR sequence *s* and the second column is for the miRNA *i* sequence *s* has been mapped to by the function *f*. Some alignment algorithms return the type of match made between a sequence *s* and the canonical sequence for miRNA *i* in the reference database. For example, miRge indicates an *exact match* between the isomiR sequence *s* and the canonical miRNA, agnostic to length, or an *isomiR* if one sequence variant exist (19). The error model we present uses *Y* and the information in the alignment data table.

### Error model

An error isomiR sequence *s*, also referred to as a technical error isomiR sequence, does not appear in the actual biological system from which the sample was taken. Rather, unwanted variation in the sequencing process resulted in errors along the read of the biological isomiR sequence *s*^′^, producing sequence *s*. Suppose we have an error isomiR sequence *s* mapping to miRNA *i* in sample *j* of a miRNA profiling experiment with associated read count *y*_*sj*_. Also mapping to miRNA *i* in sample *j* is a true biological isomiR sequence, *s*^′^, with associated read count *y*_*s*′*j*_ . The error model, including parameter estimation and hypothesis testing, relies on a pairwise alignment between the biological isomiR *s*^′^ and the sequence *s*. Here, we use pairwise alignment to mean the results of using the Needleman-Wunsch algorithm to obtain a global alignment of two nucleotide sequences (30).

Certain aspects of our error model are inspired by the DADA2 model for denoising amplicon sequencing variants (17). First, we assume that sequence read errors occur independently within a read of an isomiR sequence and between reads. From these assumptions, it follows that the probability of producing a read of *s* from *s*^′^ due to technical errors can be expressed as the product of the probabilities of the individual nucleotide transitions along the length of the pairwise alignment of *s* to *s*^′^. Let *λ*_*s*′*s*_ denote the probability of producing a read of the technical error sequence *s* from the sequence *s*^′^. Let 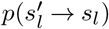 denote the probability of the nucleotide at position *l* in *s*^′^ of the pairwise alignment being read as the nucleotide at position *l* in *s* of the pairwise alignment. Finally, let *L* be the length of the pairwise alignment between *s* and *s*^′^. Then, from our assumptions, we have:

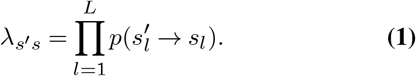

The transition probability of one character to another character is defined by the identity of those characters. We refer to these individual characters of the pairwise alignment between the sequences *s*^′^ and *s* as characters and not just nucleotides because we allow these characters to be either nucleotides or a gap: {*A, C, G, T*, −} . This allows us to handle nucleotide-to-nucleotide and gap-to-nucleotide or nucleotide-to-gap transitions simultaneously.

Each of these different types of transitions that we allow represents a different type of isomiR. Nucleotide-to-nucleotide transitions represent substitution errors. A pairwise alignment between *s* and *s*^′^ containing only nucleotide-to-nucleotide transitions indicates that *s* is a sequence variant isomiR of *s*^′^. Similarly, gap-to-nucleotide transitions represent addition errors and nucleotide-to-gap transitions represent deletion errors. A pairwise alignment between *s* and *s*^′^ that contains only gap-to-nucleotide or nucleotide-to-gap transitions indicates that *s* is a length variant isomiR of *s*^′^. Length variants, particularly at the 3^′^ end, are common in miRNA-seq because of 5^′^/3^′^ cleavage heterogeneity and 3^′^tailing (20); in microbiome amplicon sequencing, conserved primers, quality trimming, and paired-end merging produce near-uniform amplicon lengths (17). Finally, a sequence *s* with all three types of transitions is a mixed type isomiR of *s*^′^. We organize these transition probabilities in a matrix, Θ. The transition probabilities can either be prespecified or estimated directly from the data. Details regarding the methodology used for parameter estimation are in the Supplemental Materials.

Next, denote a random variable *Y*_*sj*_ representing the read count of error sequence *s* in sample *j*. Let *N*_*s*′*j*_ represent the true abundance of the biological isomiR sequence *s*^′^ in sample *j*. The value *N*_*s*′*j*_ is a latent random variable. We use *n*_*s*′*j*_ to denote its observed read count:

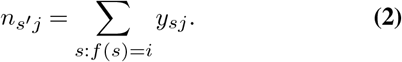

In each sample *j*, each molecule of sequence *s*^′^ is selected for sequencing with probability *p*_*s*′*j*_ . Thus, the number of observed reads originating from *s*^′^ follows

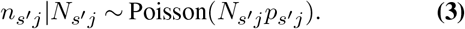

Conditional on originating from *s*^′^, a read is observed as error sequence *s* with probability *λ*_*s*′*s*_, determined by the Error model Eq Eq. (1). By the Poisson thinning property, the count of reads assigned to *s* follows:

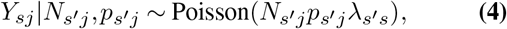

or equivalently, conditioning on *n*_*s*′*j*_ :

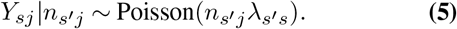

We therefore view the sequencing process for *s* as arising from two Poisson thinnings: 1) selection of reads *n*_*s*′*j*_ from the true count *N*_*s*′*j*_ based on sampling probability *p*_*s*′*j*_, and 2) misassignment of these reads to error sequence *s* with probability *λ*_*s*′*s*_.

### Hypothesis testing

We use the error model described above to test the null hypothesis *H*_0_ that sequence *s* mapping to miRNA *i* is an error isomiR sequence of biological isomiR sequence *s*^′^. Specifically, we compare the observed read count of sequence *s* in sample *j* to what we would expect under the error model from Equation 5. If *y*_*sj*_ is much more abundant than what we would expect under the error model, we reject the null hypothesis and infer that *s* is another biological isomiR sequence of miRNA *i*. We formalize the idea of being “much more abundant than what we would expect” by borrowing the concept of an abundance p-value from the DADA2 algorithm (17). The abundance p-value is defined mathematically as:

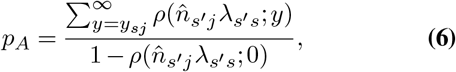

where *ρ* represents the Poisson density function. The abundance p-value is the probability of observing a read count for sequence *s* in sample *j* as extreme or more extreme than *y*_*sj*_, assuming *s* is an error isomiR sequence of biological isomiR sequence *s*^′^. The decision criteria for rejecting the null hypothesis compares the abundance p-value, *p*_*A*_, to a user-defined threshold, Ω_*A*_. Ω_*A*_ functions similarly to the *α*−level in a traditional, frequentist hypothesis testing framework and is commonly set to 0.05.

In practice, we calculate an abundance p-value for each isomiR sequence *s* such that *f*(*s*) = *i* and compare it to Ω_*A*_. To control the number of false discoveries we make, we correct the abundance p-values for multiple testing using the Benjamini-Hochberg procedure (31). We will use 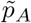 to represent the Benjamini-Hochberg adjusted abundance p-value.

### PARiS: Probabilistic Assignment and Repartitioning of isomiR Sequences

The PARiS algorithm takes, as input, a *K* ×*J* -dimensional matrix of isomiR-level read counts, *Y* . The first step of the PARiS algorithm is to estimate the transition probabilities stored in Θ using the methods for parameter estimation outlined in the Supplemental Materials. PARiS is an iterative algorithm that partitions the set of sequences 𝒮_*ij*_ mapping to miRNA *i* in sample *j* through a sequential refinement process. We drop the notation for *i* and *j* in the following. We define the initial partition by,

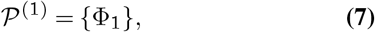

where Φ_1_ = 𝒮 for some miRNA *i*.

We use *d ∈* {1, 2, …} to index iterations of the PARiS algorithm. At iteration *d*, let 𝒫^(d)^ = {Φ_1_, …, Φ_*d*_} be a partition of 𝒮 into *d* disjoint (nonempty) subsets, i.e.,

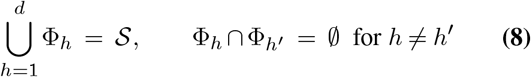

Each block Φ_*h*_ has a designated center sequence *Φ*_*h*_, which represents a biological isomiR.

At iteration (*d* + 1), we obtain a finer partition 𝒫^(*d*+1)^ by splitting the block Φ_*d*_. The steps are as follows: i) apply the error model from Equation 5 to each *s* ∈ Φ_*d*_, taking *s*^′^ = *ϕ*_*d*_ to compute the abundance p-value *p*_*A*_(*s*^′^, *s*); ii) after computing *p*_*A*_(*s*^′^, *s*) ∀*s* ∈ Φ_*d*_, we adjust for multiple testing using the Benjamini-Hochberg procedure (31) to generate 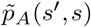 ; iii) we compare 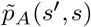 to Ω_*A*_, the user-defined significance threshold; iv) we identify a new center sequence, *Φ*_*d*+1_, by

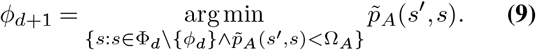

After identifying *ϕ*_*d*+1_, the set Φ_*d*+1_ is given by

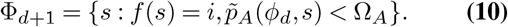

This iterative process proceeds in a waterfall-like manner, where sequences “flow” from Φ_1_ to Φ_2_, and so on, until no additional subsets can be formed or a pre-specified maximum number of iterations is reached.

After each *s* has been assigned, we have defined a partition of isomiR sequences mapping to miRNA *i* in sample *j*. We will refer to 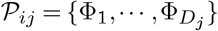 as the sample-level partition of miRNA *i* in sample *j* or 𝒫_*j*_, dropping dependence on miRNA *i* when it is clear from the context.

### Forming a consensus partition and denoising counts

After obtaining per–sample partitions for miRNA *i*, we consolidate them into a single, coherent partition by first defining a consensus set of center sequences. Let 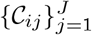 denote the sets of center sequences identified for miRNA *i* in each of the *J* samples. We define the consensus set of center sequences for miRNA *i* as 𝒞_*i*_.

Intuitively, 𝒞_*i*_ contains the center sequences that appear in a sufficient proportion of samples. Two extremes illustrate this idea:

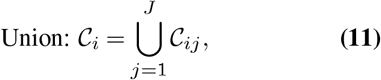

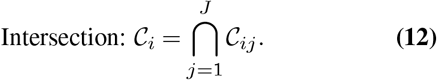

Taking the union is permissive and risks false positives (including error isomiRs as centers), while taking the intersection is stringent and risks false negatives (excluding true biological isomiRs).

To balance these extremes, we introduce a prevalence threshold β ∈ [1*/J*, 1]: a candidate center is retained in 𝒞_*i*_ if it appears in at least a *β* fraction of samples. Setting *β* = 1*/J* yields the union, whereas *β* = 1 yields the intersection. The resulting consensus set 𝒞_*i*_ defines the blocks of the consensus partition, which we then use to denoise counts across samples.

Upon identification of the consensus center set 𝒞_*i*_, we reassign each non-center sequence *s* ∈ 𝒮_*i*_ \ 𝒞_*i*_ as follows. For every center *s*^′^ ∈ 𝒞_*i*_, compute *λ*_*s*′*s*_ as in Eq. 1, and assign *s* to the block whose center attains the largest *λ*_*s*′*s*_. The total number of subsets for *i* is given by the cardinality of 𝒞_*i*_: *D*_*i*_ = |𝒞_*i*_|.

After constructing the consensus partition 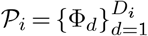, we denoise counts by aggregating reads within blocks. Let *ϕ*_*d*_ ∈ 𝒞_*i*_ denote the center of block Φ_*d*_. For sample *j*, the denoised count for putative biological isomiR sequence *ϕ*_*d*_ is

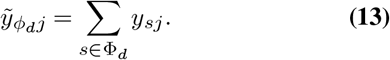

The inferred composition for miRNA *i* is given by its consensus centers 𝒞_*i*_, with denoised abundances 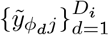 across samples.

### Simulating data from the error distribution

We begin by benchmarking the ability of PARiS to identify and remove technical error isomiR sequences without introducing new errors. To do this, we require a dataset where, for each isomiR sequence *s* we know whether *s* is a biological isomiR or an error isomiR. We generated our own miRNA sequencing data with associated read counts and ground truth error isomiR sequences from an experimental benchmark dataset sequencing mouse miRNAs. The experimental benchmark dataset we used consisted of 421,096 sequences mapping to 759 distinct miRNAs after alignment to miRGeneDB (8) using sRN-Abench (21). We filtered out lowly expressed miRNAs with median counts per million less than 5. After filtering 435 miRNAs remained. For each miRNA, we identified *s*^′^, the most abundant sequence mapping to the miRNA across all samples. Then, for each miRNA, we simulated making additions, deletions, or substitutions to the string representing *s*^′^ to produce isomiR sequences from the center sequence. We then simulated read counts from the error model given in Equation 5 to generate read counts of the simulated isomiR sequences. The details of the data simulation can be found in the Supplemental materials.

We varied the number of true biological isomiR sequences mapping to each miRNA by selecting the value from the set {1, 4, 9}. Then, for a given true number of biological isomiR sequences, we repeated the process of generating simulated isomiR sequences and simulating read counts from the error model to generate a simulated dataset 10 times.

### Simulated monocyte datasets

To demonstrate that PARiS improves miRNA-level differential expression analysis, we require a dataset with several key characteristics. First, the dataset must have isomiR sequence read counts. Second, the dataset must have ground truth values for the amount of differential expression for each of the miRNA in the dataset. Differential expression is typically measured by the log fold change (logFC) between experimental conditions, thus we must either know the true logFC for each miRNA or be able to compute it.

To generate a dataset that meets these requirements, we use the same process used by Baran et al. to artificially inject differential expression signal into a real dataset (15). The initial dataset consisted of 39 monocyte samples. After filtering for lowly expressed miRNAs and removing sequences that were not an exact match to the reference database, the initial dataset consists of 122 miRNAs mapping to 3538 unique sequences. From this initial dataset, 50 simulated datasets were created using the following process: The 39 samples were split into two groups (A and B) randomly to mimic a simple experimental design (15). From the 122 miRNAs, 20 were selected to be overexpressed in group *A* at random and 20 were selected to be overexpressed in group *B* at random. The isomiR-level read counts of the miRNAs selected to be overexpressed in each group were multiplied by values sampled from a truncated normal distribution with a mean of 2 and a standard deviation of 1. More details about this process can be found in Baran *et al*. (15). The simulation process was repeated to generate 50 synthetic datasets, with known logFC values for each miRNA and each isomiR sequence in the dataset. We do not observe whether each isomiR sequence is a biological isomiR sequence or an error isomiR sequence.

We applied different analytical pipelines, consisting of a denoising method selected from the set of {PARiS, miREC, Aggregation, or None} and an appropriate method for estimating miRNA-level differential expression given the resolution of the denoised data. For the analytical pipeline using PARiS, we used Ω_*A*_ = 0.05 as the threshold for identifying significant abundance p-values. We varied the *β* value for creating consensus sets of center sequences for each miRNA by selecting *β* from {0.025, 0.05, 0.10, 0.20, 0.40, 0.60, 0.80, 1.00}. PARiS, and miREC produce isomiR-level denoised data, and the raw data (the None denoising option) is also at the isomiR-level. We estimated miRNA-level differential expression using the denoised data from each of these methods using miRglmm, a generalized linear mixed effects modeling framework (15). The aggregation method produces denoised data at the miRNA-level. We estimated miRNA-level differential expression from the aggregated data using DESeq2 (14) and edgeR (13). All together, we applied 11 different analytical pipelines to each of the 50 simulated datasets.

### ERCC benchmark dataset

We evaluated the performance of PARiS in an analytical pipeline for miRNA differential expression analysis on a synthetic, experimental benchmark dataset. Because of the synthetic nature of the dataset, there is no biological variation present. We used the benchmark dataset from the Extracellular RNA Communication Consortium (ERCC) (22). The ERCC dataset is sRNA-seq on ratio-metric pools (A and B) of synthesized small RNAs in ratios from 10:1 to 1:10. The result is an isomiR sequence expression dataset with 15 levels of ground truth differential expression, ranging from log(0.1) = −2.3 to log(10) = 2.3. The dataset contains 286 human miRNAs mapping to 8001 sequences after removing sequences that are ≥ 40 nucleotides or *<* 16 nucleotides.

We denoise the ERCC dataset using a similar set of methods that were applied to the simulated monocyte datasets. The denoising methods we applied to the ERCC dataset are as follows: miREC, aggregation, and PARiS + *β* ∈ { 0.05, 0.1, 0.15, 0.2, 0.25, 0.3, 0.35, 0.4, 0.45, 0.5, 0.55, 0.6, 0.65, 0.7, 0.75, 0.8, 0.85, 0.9, 0.95, 1} . For each application of PARiS, we set Ω_*A*_ = 0.05. Together, this gives a total of 22 different analytical pipelines. Following denoising in each pipeline, we estimated differential expression at the miRNA-level using the appropriate method given the resolution of the denoised data. For the miREC-corrected and PARiS-denoised data, we estimated miRNA-level differential expression using miRglmm (15). For the aggregated data, we estimate miRNA-level differential expression using DESeq2 (14) and edgeR (13).

### Comparison of colorectal adenocarcinoma cell lines

We applied PARiS as the denoising step in a miRNA differential expression analysis pipeline applied to a true, experimental dataset. The sRNA-seq data (N=9) are from 3 colerectal adenocarcinoma cell lines (DLD-1, DKO-1, and DKS-8), which vary by KRAS status (23). The raw sequencing data consists of 5223 unique sequences mapping to 143 miRNAs after filtering out lowly expressed miRNAs with reads per million < 5 and keeping only sequences that exactly aligned to the reference database. We applied PARiS with *β* = 0.4 and Ω_*A*_ = 0.05 and then estimated miRNA-level differential expression using miRglmm (15). We also aggregated isomiR-level counts to the miRNA-level and estimated differential expression using DESeq2 for comparison purposes (14). We identified miRNAs as differentially expressed using a significance level of *α* = 0.05. For miRNAs identified as differentially expressed by DESeq2 and not miRglmm, we tested for differential isomiR usage using a likelihood ratio test.

### Performance evaluation metrics

#### Simulated null data

For each miRNA in the datasets simulated from the error model, we know whether each sequence *s* is a biological isomiR sequence or an error isomiR sequence. To evaluate the performance of the error model on the simulated data, we report the false positive rate (FPR) and true positive rate (TPR):

- False positive rate: 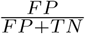, shows the proportion of error isomiR sequences misidentified as biological isomiR sequences by the algorithm.
- True positive rate: 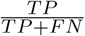, shows the proportion of biological isomiR sequences correctly identified as biological isomiR sequences by the algorithm.

where true positives *TP* are correctly identified biological isomiR sequences, *FP* are error isomiR sequences that have been labeled biological isomiR sequences by the algorithm, *TN* are error isomiR sequencces that have been labeled error isomiR sequences by the algorithm, and *FN* are biological isomiR sequences that have been labeled as error isomiR sequences by the algorithm. Since we applied PARiS to 10 simulated datasets for each of the simulation settings, we report the distribution of the FP and TP rates over the simulated datasets.

#### Simulated datasets and experimental benchmark dataset with ground truth log-fold change

For both the 50 simulated monocyte datasets and the ERCC dataset, we evaluate the performance of a method on a given dataset with the mean squared error (MSE) of the log fold change estimates and with the coverage proportion of the estimated 95% confidence intervals. We use the definitions for both terms commonly given in the literature.

- Mean squared error: 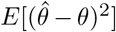, where 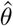 are the estimated log fold change values for each miRNA and *θ* are the induced log fold change effect, either by manipulating the count data or creating experimental pools with known concentration ratios.
- Coverage proportion: Proportion of true log fold changes *θ* that fall within the estimated 95% confidence interval.

### Software, data structures, outputs, and reproducibility

The PARiS algorithm is written in the R programming language. We use the ‘SummarizedExperiment’ package in the R programming language to organize the count data, *Y*, the data table with the information from the alignment defined by function *f*, and additional sample-level covariate information into a single object. In the language of the SummarizedExperiment package, the count data *Y* is an assay. The data table with the additional data produced by the alignment is the ‘rowData’ and the data table with the sample-level covariate information is the ‘colData’. Many of the R packages created to perform differential expression analysis, of either miRNA data or bulk RNA-seq data, expect a Summarized-Experiment object with these elements as input. Therefore, PARiS takes a SummarizedExperiment object as input and returns a denoised SummarizedExperiment object as output.

## Results

### The error model in PARiS successfully identifies technical length variant and sequence variant error isomiR sequences in simulated data

We begin with the results from using the proposed error model that PARiS is based on to identify technical length variant error isomiR sequences. We allow the true number of biological isomiR sequences per miRNA to take on a different value from the set {1, 4, 9} . For each miRNA in the dataset, we calculated the false positive rate according to the definition provided in the previous section. When there was 1 true biological isomiR per miRNA, the average false positive rate is approximately 0.015 (Figure 1, A, top). When there are 4 true biological isomiRs per miRNA, the average false positive rate was 0.09 (Figure 1, B, middle). Finally, for the scenario where there are 9 true biological isomiR sequences per miRNA, the average false positive rate was 0.10 (Figure 1, A, bottom).

**Fig. 1.**
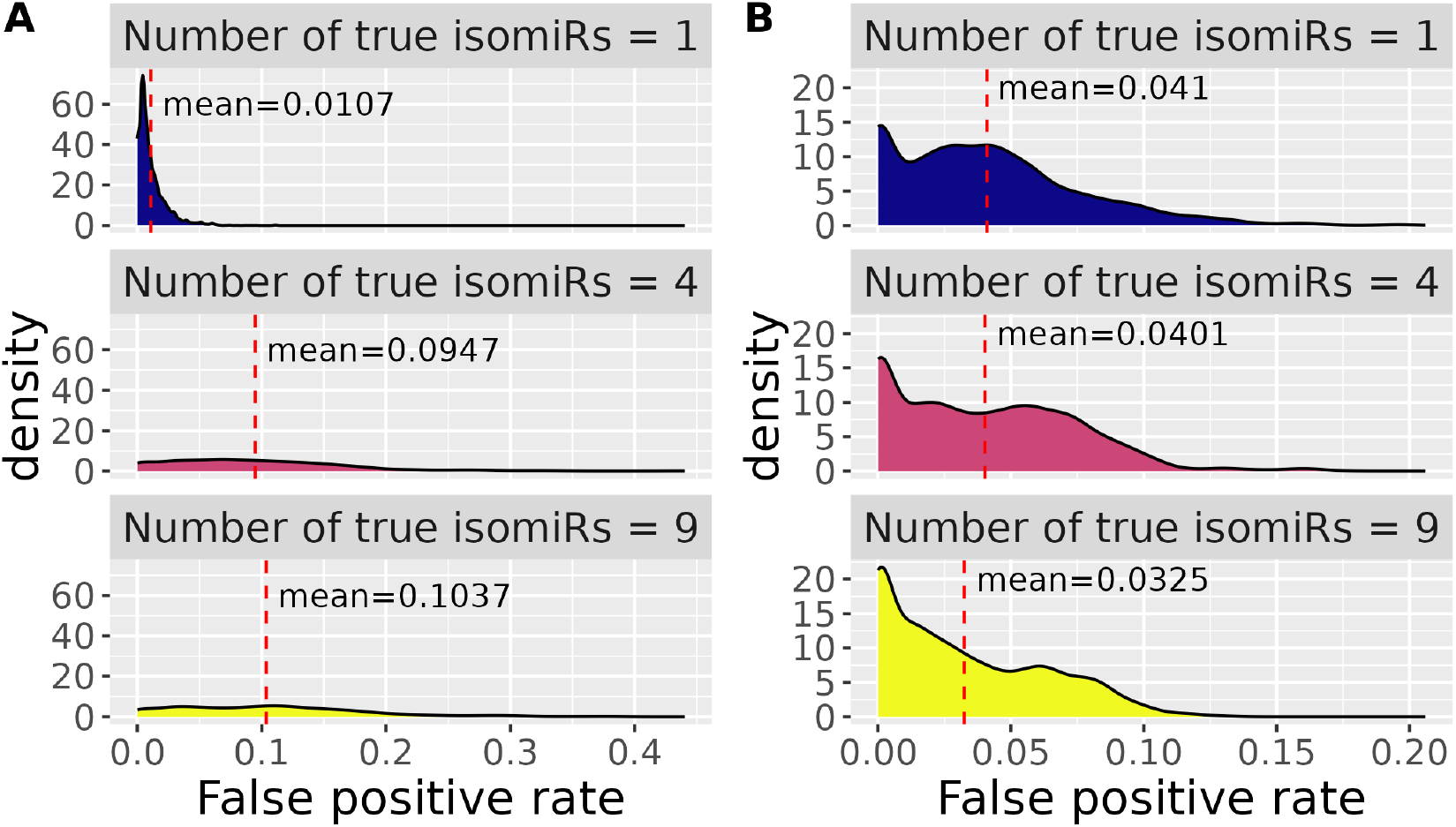
Results of using the proposed error model to identify technical error isomiR sequences in simulated datasets with varying numbers of true biological isomiR sequences. Panel A shows the distribution of the false positive rates, over 10 simulations, by the number of true isomiRs per miRNA in the simulated datasets with only length variant isomiR sequences. Panel B shows the distribution of the false positive rates, over 10 simulations, by the number of true isomiRs per miRNA in the simulated datasets with only sequence variant isomiR sequences. Each row corresponds to a specific true number of isomiRs per miRNA from the set {1, 4, 9}. The average false positive rate over the 10 simulations for a given simulation setting is represented on the plot with a dashed, vertical line in red.

Length variant isomiRs are not the only type of isomiR sequence potentially present in the data. Sequence variant isomiRs are also potentially present, and it is important that the error model in PARiS can handle these as well. We simulated sequence variant isomiRs from the data, again letting the true number of biological isomiR sequences take on different values from the set {1, 4, 9}. For each miRNA, we report the false positive rate and true positive rate of identifying technical error sequence variant isomiRs from the data. When there is only 1 true biological isomiR per miRNA, the average false positive rate was approximately 0.04 (Figure 1, B, top). When there are 4 true biological isomiRs per miRNA, the average false positive rate was still approximately 0.04 (Figure 1, B, middle). Finally, when there are 9 true biological isomiRs per miRNA, the average false positive rate was 0.03 (Figure 1, B, bottom).

### PARiS decreases the MSE and increases the coverage proportion of miRNA-level log fold change estimates using isomiR-level expression data in simulated data

We begin by examining the MSE of the different analytical pipelines used to denoise the data and estimate miRNA-level differential expression in the set of 50 simulated monocyte datasets. A lower MSE indicates that the analytical pipeline generates logFC estimates that are, on average, closer to the ground-truth logFC values. Across the 50 simulated datasets, miRglmm models fit to isomiR-level count data denoised with PARiS, with Ω_*A*_ = 0.05 and *β* = 0.40 had the lowest mean MSE (MSE=0.0162) of all the analytical pipelines assessed (Table 1). Surprisingly, the average MSE over the 50 simulated datasets is actually greater for the miRglmm models fit to the miREC-corrected data (mean=0.034) than for the miRglmm models fit to the raw data (mean=0.033).

**Table 1.** Performance results of different analytical pipelines across 50 simulated monocyte miRNA expression datasets in terms of the mean squared error (MSE) of the miRNA-level log fold change (logFC) estimates. The smaller the MSE, the closer, on average, the logFC estimates from the analytical pipeline are to the true values. The method column refers to the type of method used either to denoise the data (PARiS, miREC, None), or, in the case where the sequence-level counts were aggregated to the miRNA-level, the method used to estimate miRNA-level differential expression (DESeq2, edgeR). For estimating miRNA-level differential expression in sequence-level datasets, we used the miRglmm modeling framework. For the analytical pipelines that used PARiS, the user must select values for Ω_*A*_ and *β*. We used Ω_*A*_ = 0.05 for all applications of the PARiS algorithm. We varied *β*, the value used for generating consensus sets of center sequences from 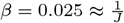 to *β* = 1 to evaluate how performance changes as *β* changes.

| Method | Average MSE<br>($10^{-3}$) | Min MSE<br>($10^{-3}$) | Max MSE<br>($10^{-3}$) | # of times mini-<br>mized MSE |
| --- | --- | --- | --- | --- |
| PARiS, $\beta = 0.025$ | 52.19 | 16.92 | 93.75 | 0.00 |
| PARiS, $\beta = 0.05$ | 21.35 | 9.62 | 43.68 | 1.00 |
| PARiS, $\beta = 0.1$ | 18.02 | 7.71 | 39.90 | 3.00 |
| PARiS, $\beta = 0.20$ | 16.75 | 7.19 | 37.12 | 7.00 |
| PARiS, $\beta = 0.40$ | 16.23 | 6.70 | 33.81 | 12.00 |
| PARiS, $\beta = 0.60$ | 16.24 | 6.51 | 32.74 | 10.00 |
| PARiS, $\beta = 0.80$ | 16.57 | 6.99 | 30.40 | 9.00 |
| PARiS, $\beta = 1.00$ | 17.65 | 10.12 | 30.01 | 4.00 |
| miREC | 34.40 | 21.54 | 68.42 | 0.00 |
| None | 33.31 | 17.74 | 65.33 | 0.00 |
| DESeq2 | 22.50 | 15.34 | 31.12 | 4.00 |
| edgeR | 24.23 | 15.32 | 34.78 | 0.00 |

The analytical pipeline that denoised the data with PARiS, with Ω_*A*_ = 0.05 and *β* = 0.025, and estimated miRNA-level differential expression using miRglmm generated logFC estimates with the highest average MSE. The analytical pipelines that used PARiS to denoise the data with Ω_*A*_ = 0.05 and any other value of *β*, then estimated miRNA-level differential expression using miRglmm, generated logFC estimates with smaller average MSE than the data denoised with any other method.

In addition to MSE, we also evaluated the analytical pipelines in terms of the coverage proportion of the estimated 95% confidence intervals. Estimating confidence intervals requires point estimates for the parameter of interest and an estimate of the standard error. Because edgeR does not provide the user with standard error estimates, we only report performance in terms of coverage proportion for the analytical pipelines that estimated differential expression using either miRglmm or DESeq2.

Of all analytical pipelines used to estimate differential expression, the analytical pipelines that denoised the data with PARiS, with Ω_*A*_ = 0.05 and *β* ∈ {0.05, 0.1, 0.2, 0.4, 0.6}, then estimated differential expression using miRglmm estimated 95% confidence intervals with the greatest average coverage proportion (mean coverage proportion = 0.90) (Table 2). The next highest coverage proportion came from the analytical pipeline that denoised the data using PARiS, with Ω_*A*_ = 0.05 and *β* ∈ {0.8, 1}, then estimated differential expression with miRglmm. Regardless of the *β* value used, any analytical pipeline that denoised the data with PARiS and estimated differential expression with miRglmm estimated 95% confidence intervals with higher average coverage proportion than competing methods. Any analytical pipeline using PARiS, regardless of *β* value used, also estimated 95% confidence intervals with greater minimum coverage proportion and maximum coverage proportion over the 50 simulations.

**Table 2.** Performance results of different analytical pipelines across 50 simulated monocyte miRNA expression datasets in terms of the coverage proportion of the estimated 95% confidence intervals. The greater the coverage proportion, the better the performance. The methods used are the same methods used in Table 1. If the “denoising” step of the analytical pipeline returns sequence-level counts (PARiS, miREC, None), the methods column lists the type of denoising used. For applications of PARiS, this includes listing the *β* value used to generate consensus sets of center sequences. Ω_*A*_ = 0.05 was used for all applications of PARiS. For analytical pipelines that aggregate sequence-level counts to the miRNA-level, the method column describes the method used for estimating differential expression (DESeq2). There are no results reported for the analytical pipeline that aggregates sequence-level counts to the miRNA-level and estimates miRNA-level differential expression using edgeR because edgeR does not report standard errors of the logFC estimates. For analytical pipelines where the denoising step returned sequence-level counts (PARiS, miREC, None), miRNA-level differential expression was estimated using the miRglmm modeling framework.

| Method | Average coverage proportion | Min coverage proportion | Max coverage proportion |
| --- | --- | --- | --- |
| PARiS, $\beta = 0.025$ | 0.88 | 0.71 | 0.98 |
| PARiS, $\beta = 0.05$ | 0.90 | 0.74 | 0.98 |
| PARiS, $\beta = 0.10$ | 0.90 | 0.74 | 1.00 |
| PARiS, $\beta = 0.20$ | 0.90 | 0.72 | 1.00 |
| PARiS, $\beta = 0.40$ | 0.90 | 0.74 | 1.00 |
| PARiS, $\beta = 0.60$ | 0.90 | 0.74 | 1.00 |
| PARiS, $\beta = 0.80$ | 0.89 | 0.73 | 0.98 |
| PARiS, $\beta = 1.0$ | 0.89 | 0.71 | 0.97 |
| miREC | 0.86 | 0.67 | 0.96 |
| None | 0.86 | 0.70 | 0.95 |
| DESeq2 | 0.81 | 0.71 | 0.88 |

In addition to evaluating the performance of the different analytical pipelines in terms of MSE and coverage proportion on all miRNAs, we also assessed each method’s performance within each of the truth groups in the monocyte simulations. Each dataset in the set of 50 simulated monocyte datasets contains 3 truth groups. The first group is the group of miRNAs with induced positive change from group *B* to group *A* (logFC equal to log(2). The second group is the group of miRNAs with induced negative change from group *B* to group *A* (logFC equal to log(0.5). The third and final group is the group of miRNAs with no induced change from group *B* to group *A* (logFC equal to log(1). The distribution of the MSEs across the 50 simulations for each analytical pipeline, within truth group, is shown in Figure 2 (A). The distribution of the coverage proportion across the 50 simulations for each analytical pipeline, within truth group, is shown in Figure 2 (B). Again, no coverage proportion results for the analytical pipeline that aggregates sequence-level read counts to the miRNA-level and uses edgeR to estimate differential expression are reported. This is because edgeR does not report standard errors of the logFC estimates.

**Fig. 2.**
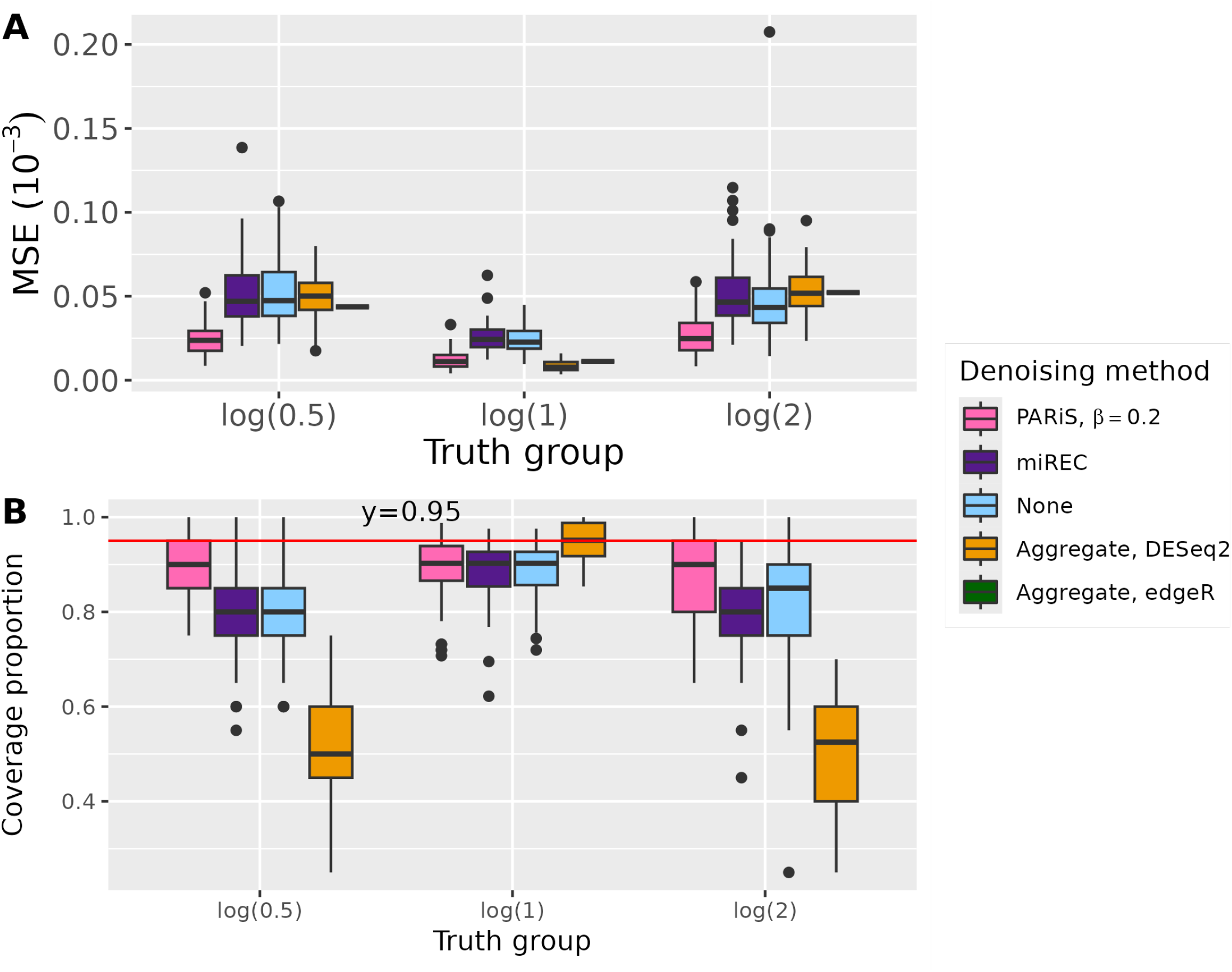
(A) Results of using different analytical pipelines to denoise the sequence-level reads and estimate miRNA-level differential expression in 50 simulated monocyte datasets by true log fold change (truth group). Each boxplot represents the mean squared error (MSE) (y-axis) of the log fold change (logFC) estimates by truth group (x-axis) across the 50 simulated datasets for each approach to denoising. The fill color of the boxplot represents the different analytical pipelines. For denoising done with PARiS, we used Ω_*A*_ = 0.05 and *β* = 0.2. For the aggregation approach, we label the boxplot according to the differential expression estimation method used, DESeq2 or edgeR. In the PARiS denoised datasets, the miREC denoised datasets, and the raw data, miRNA-level differential expression was estimated from sequence-level data using miRglmm. (B) Each boxplot represents the coverage proportion of the estimated 95% confidence intervals (y-axis) of the logFC estimates by truth group (x-axis) across the 50 simulated datasets for each approach to denoising. The color of each boxplot corresponds to method in the same manner as panel A. In panel B, for the aggregated dataset, we only report results for DE estimation with DESeq2 because edgeR does not provide standard error estimates. We indicate the nominal 95% level with a horizontal red line.

In the groups with an induced effect, using an analytical pipeline that uses miRglmm to estimate differential expression after denoising with PARiS, with Ω_*A*_ = 0.05, regardless of the selected *β* value, produces logFC estimates with MSE less than or equal to the MSE of the logFC estimates prodced by denoising with competing methods (Figure 2, A). In the group with no logFC, the analytical pipelines that use aggregation techniques and estimate differential expression using either DESeq2 or edgeR minimizes MSE (Figure 2, A). However, all analytical pipelines achieve comparably low MSE in the truth group with no induced effect, except for the piepline that uses PARiS with Ω_*A*_ = 0.05 and *β* = 0.025.

Similarly, in groups with an induced effect, using an analytical pipeline that uses miRglmm to estimate differential expression after denoising with PARiS, with Ω_*A*_ = 0.05, regardless of the selected *β* value, estimates 95% confidence intervals with coverage proportions greater than or equal to the coverage proportions of the estimated 95% confidence intervals from competing methods (Figure 2, (B)). In the truth group with no induced effect, the analytical pipeline that aggregates sequence-level read counts to the miRNA-level and estimates differential expression using DESeq2 estimates 95% confidence intervals with the highest coverage proportion. While the DESeq2 analytical pipeline achieves the highest coverage proportion, all competing methods achieve comparably high coverage of the estimated 95% confidence intervals in the truth group with no induced change.

### PARiS identifies and removes isomiR sequences miREC misses, thus improving miRNA-level log fold change estimates of isomiR-level expression data when there is no biological sequence variability

First, we look at how the distribution of the number of isomiRs per miRNA changes in the PARiS-denoised data vs. the miREC-corrected data as we change the value of *β* used to generate consensus sets of center sequences. In the miREC-treated data, the average number of isomiRS per miRNA is 0. Given the synthetic nature of the ERCC data, the true number of isomiRs per miRNA is 1. Denoising the data with Ω_*A*_ = 0.05 and any *β* value moves the distribution of isomiRs per miRNA away from the distribution of the raw data and towards the true value (Figure 3, panel A).

**Fig. 3.**
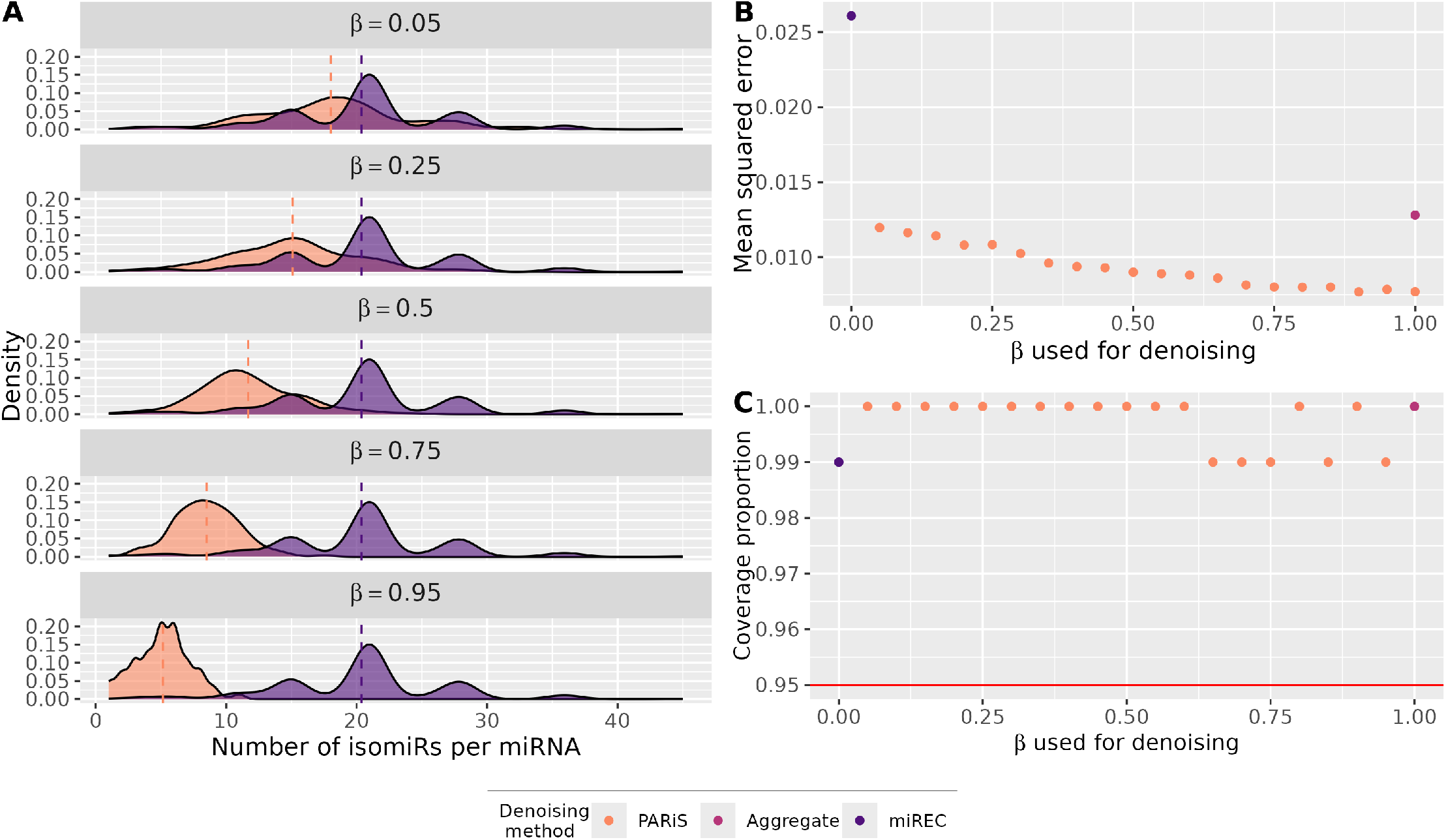
(A) Distributions of the number of isomiRs per miRNA in the raw data and the data denoised with PARiS with Ω_*A*_ = 0.05, and the *β* value ranging from 0.05 to 0.95. Given the synthetic nature of the dataset, we know that the ground truth for each miRNA is 1 true biological isomiR per miRNA. As *β* increases towards 1, the average number of isomiRs per miRNA in the denoised data moves closer to the ground truth. (B) MSE for MIREC denoised data, non-denoised aggregate data, and PARiS denoised data with different *β* values. (C) Coverage proportions for MIREC denoised data, non-denoised aggregate data, and PARiS denoised data with different *β* values.

Next, we evaluated the different analytical pipelines in terms of the MSE of the logFC estimates each pipeline produced. Recall that the smaller MSE, the better the method is performing because the closer, on average, the estimates are to the true values. Using an analytical pipeline that uses PARiS to denoise the data, with Ω_*A*_ = 0.05, regardless of the *β* value, produces miRNA-level logFC estimates with smaller MSE than those produced by the other analytical pipelines (Figure 3, panel B). Furthermore, as *β* increases from 0.025 towards 1, the MSE of the logFC estimates decreases. The analytical pipeline that denoised the data with miREC and estimated differential expression with miRglmm produced miRNA-level logFC estimates with the largest MSE. Finally, we used the coverage proportions of the estimated 95% confidence intervals from the different analytical pipelines to evaluate them. The definition of the coverage proportion is the same as the definition used to evaluate the different analytical pipelines on the set of simulated monocyte datasets. That is, the coverage proportion is the proportion of miRNAs whose true logFC value falls within the estimated 95% confidence interval. The greater the coverage proportion, the better the method is performing. All of the analytical pipelines estimated 95% confidence intervals with coverage proportions at or above the nominal 0.95 level (Figure 3, panel c). Although there is variability in the coverage proportion of the 95% confidence intervals with respect to the *β* value used in PARiS to denoise the data, it is minimal. Using an analytical pipeline that denoises the data with PARiS, with Ω_*A*_ = 0.05 and any *β* value, estimates 95% confidence intervals with coverage proportions greater than the nominal 0.95 level. This indicates that denoising with PARiS improves point estimates without sacrificing the standard error estimates of the point estimates.

### Using PARiS as a step in analytical pipeline applied to a real, experimental dataset may prevent researchers from mistaking differential isomiR usage as differential expression

We performed multiple pairwise comparisons of three colorectal adenocarcinoma cell lines. The first comparison we made compared the isomiR expression profiles of the DKO-1 and DKS-8 cell lines and the second comparison we made compared the isomiR expression profiles of the DKO-1 and DLD-1 cell lines. We used two different analytical pipelines for the differential expression analysis, one that denoised the data with PARiS, using Ω_*A*_ = 0.05 and *β* = 0.30 and estimated differential expression using miRglmm. The other analytical pipeline aggregated the isomiR-level data to the miRNA-level and estimated differential expression using DESeq2. The analytical pipeline using PARiS and miRglmm found 53 miRNAs to be differentially expressed between the DKS-8 and DKO-1 cell lines (Figure 4, panel A). The analytical pipeline using aggregation and DESeq2 found 34 miRNAs to be differentially expressed between the DKS-8 and DKO-1 cell lines. The two pieplines shared 19 miRNAs in common.

**Fig. 4.**
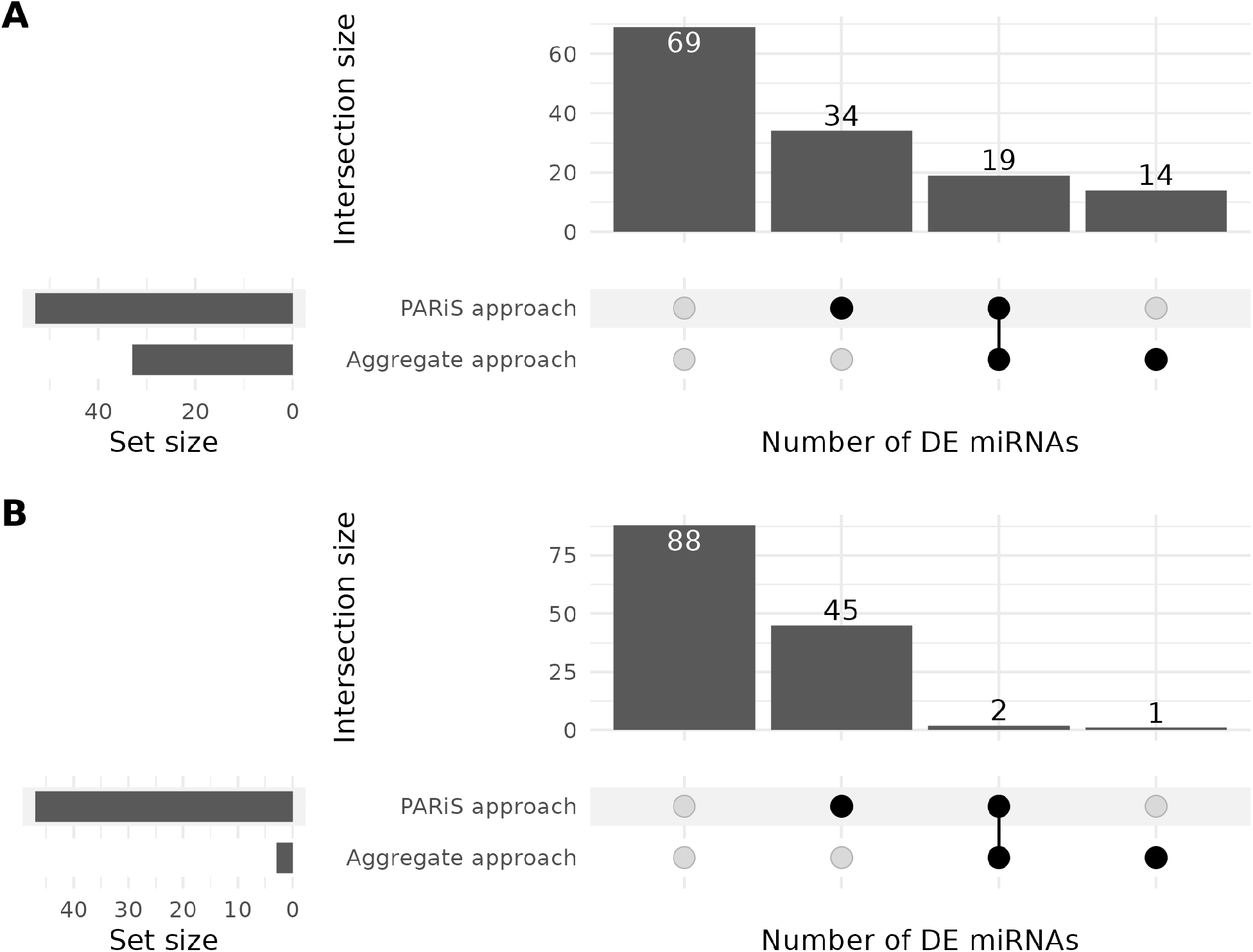
(A) Upset plot showing the number of miRNAs identified between DKS-8 and DKO-1 cell lines of colorectal adenocarcinoma cells as differentially expressed by one or more analytical pipelines applied to the data. The DKS-8 and DKO-1 cell lines differ based on the knockout status of the *Ki-RAS* alleles. Cells from the DKS-8 cell line have one wild-type *Ki-ras* allele. Cells from the DKO-1 cell line have one knocked out *Ki-RAS* allele. The first analytical pipeline aggregates sequence-level counts to the miRNA-level and estimates differential expression using DESeq2. The second analytical pipeline denoises the data using PARiS, with Ω_*A*_ = 0.05 and *β* = 0.30 and estimates differential expression using miRglmm. (B) Upset plot showing the number of miRNAs identified between DLD-1 and DKO-1 cell lines of colorectal adenocarcinoma cells as differentially expressed by one or more analytical pipelines applied to the data. Cells from the DLD-1 cell line have one wild type *Ki-RAS* allele and one mutated *Ki-RAS* allele. The analytical pipelines used in these analyses are the same as the ones used to generate the results shown in panel A.

Next, we compared miRNA expression patterns between the DLD-1 and DKO-1 cell lines. The PARiS / miRglmm identified 45 differentially expressed miRNAs between the DLD-1 and DKO-1 cell lines (Figure 4, panel B). The aggregation / DESeq2 method identified 3 miRNAs as differentially expressed. Two miRNAs were shared by both approaches.

In addition to estimating miRNA-level logFC estimates, we can also use the miRglmm modeling framework to perform a likelihood ratio test for differential isomiR usage (15). It is hypothesized that in differential isomiR usage, miRglmm will return logFC estimates that are closer to 0 than aggregation methods. In this scenario, miRglmm often finds only differential isomiR usage and does not identify miRNA-level differential expression. In our comparison of the DKO-1 and DKS-8 cell lines, we examine this hypothesis. There are 14 miRNAs differentially expressed between the DKO-1 and DKS-8 cell lines by the analytical pipeline using DESeq2 but not PARiS / miRglmm (Figure 4, panel A). Of those 14 miRNAs, the likelihood ratio test identified 6 of them as having differential isomiR usage (Table 3). Finally, of the 6 miRNAs inferred to have differential isomiR usage, 5 had smaller logFC estimates from the miRglmm modeling framework than the DESeq2 logFC estimates.

**Table 3.** miRNAs identified as differentially expressed between the DKO-1 and DKS-8 cell lines by only DESeq2 and identified as having differential isomiR usage using the likelihood ratio testing framework developed by Baran *et al*. in their miRglmm manuscript.

| miRNA |
| --- |
| hsa-miR-200a-3p |
| hsa-miR-7-5p |
| hsa-miR-221-3p |
| hsa-miR-25-5p |
| hsa-let-7i-5p |
| hsa-miR-29b-3p |

## Conclusions

There are multiple isomiR sequences mapping to a miRNA in miRNA profiling experiments, and studies have shown that these isomiRs are biologically produced and functional. However, isomiRs are not exclusively produced via biosynthesis pathways. Technical variation resulting from errors in the sequencing process of true biological isomiRs can produce isomiR sequences as well. To reduce the impact of these errors on isomiR analysis, we have developed PARiS, a method to reassign erroneous sequences to their likely source.

We demonstrate here, in both a simulation setting and on a synthetic, experimental benchmark dataset, that denoising with PARiS and estimating miRNA-level differential expression using miRglmm improves inference. The improvements in inference are reflected in both the decrease in mean squared error (MSE) of the miRNA-level logFC estimates and in the increase in the coverage proportion of the estimated 95% confidence intervals. We compare our proposed analysis pipeline to two commonly applied pipelines. The first applies miREC to the data to identify and fix error reads, then estimates miRNA-level differential expression using miRglmm. The second aggregates sequence-level counts to the miRNA-level, then estimates miRNA-level differential expression using DESeq2. Ideally, after using a denoising method (PARiS, miREC, or aggregation) we could estimate miRNA-level differential expression using the same method. However, there is currently no such method. A mixed effects modeling framework is implemented in miRglmm and requires at least 2 sequences mapping to a given miRNA to estimate random effects. On the other hand, DESeq2 requires the reads be collapsed to the miRNA-level, which is equivalent to assuming there is 1 true biological isomiR sequence per miRNA. An ideal miRNA-level differential expression modeling framework would be able to handle both of these scenarios simultaneously. Nonetheless, in our analysis of the 50 simulated monocyte datasets, we can compare the performance of miRglmm on the miREC-treated and PARiS-treated data to the performance of miRglmm on the raw data to get a sense of how denoising changes miRglmm’s performance. Assuming we are using miRglmm to estimate miRNA-level differential expression, we see that denoising with PARiS, for Ω_*A*_ = 0.05 and a wide range of *β* values improves inference over the raw data.

Finally, we apply PARiS with *β* = 0.30 and Ω_*A*_ = 0.05 to a real analysis of 3 colorectal adenocarcinoma cell lines: DLD-1, DKS-8, and DKO-1. The three cell lines differ in the knockout status of their KRAS alleles, where DKS-8 is wildtype, DLD-1 is a heterozygous KRAS mutant and DKO-1 is a homozygous KRAS mutant. Because the DLD-1 cell line is heterozygous, its expression level of KRAS would fall between the expression levels of the DKO-1 and DKS-8 cell lines. Although smaller logFCs are more difficult to detect (as expected between DLD-1 and the other cell lines), the approach of denoising with PARiS and estimating differential expression with miRglmm can identify these changes. For example, miRglmm identifies hsa-miR-200c-3p is differentially expressed between the DLD-1 and DKO-1 cell lines; DESeq2 does not make this discovery. Previous studies examining the link between miRNA expression and colorectal cancers have identified significant up-regulation of miR-200c-3p in colorectal cancer cells with a KRAS gene mutation (35). Thus, it is likely that the isomiR-based PARiS/miRglmm approach identifies more biologically relevant results than the aggregation/DESeq2 approach.

A sequence-level treatment of the data, which relies on having a high-quality, denoised version of the data, also prevents researchers from mistaking differential isomiR usage for miRNA-level differential expression. For example, in a comparison of the miRNA expression profiles between the DKS-8 and DKO-1 cell lines, there are 14 miRNAs identified as differentially expressed by DESeq2 and not miRglmm (Figure 4, 2). Baran *et al*. noted that it is possible that when there is differential isomiR usage, miRglmm will return logFC estimates that are closer to 0 than aggregation methods, not identify those miRNAs as differentially expressed, but will identify those miRNAs as having differential isomiR usage (15). Of the 14 miRNAs identified as differentially expressed, 6 have differential isomiR usage (Table 3). Of those 6, miRglmm returned logFC estimates closer to 0 for 5 of them than DESeq2. Thus, analyzing the data at the sequence-level allows researchers to improve inference both by potentially identifying differentially expressed miRNAs with logFCs of smaller magnitudes and not mistaking differential isomiR usage for differential miRNA expression.

In conclusion, we have developed PARiS, an algorithm that denoises isomiR sequencing data through the Probabilistic Assignment and Repartitioning of isomiR sequences. In data simulated from the proposed error model used by PARiS, the model performs as expected. In settings in which ground truth differential expression as measured by the logFC, is known, denoising with PARiS and estimating differential expression with miRglmm, a negative binomial mixed effects modeling framework developed for estimating miRNA-level differential expression from sequence-level data, improves inference. The improvement in inference is shown both in a decrease in MSE and an increase in the coverage proportion of the 95% confidence intervals. Furthermore, a real data analysis using PARiS improved inference by correctly identifying miRNAs with smaller magnitude logFCs as differentially expressed and not mistaking miRNAs with differential isomiR usage as differentially expressed.

## ACKNOWLEDGEMENTS

This work was supported by NIH grant R01GM139928.

## Supplementary Note 1: Transition probability matrix estimation

The error model used in the PARiS algorithm uses the probabilities of different types of transitions in the pairwise alignment between the biological isomiR sequence and error isomiR sequence as parameters. The user can provide the PARiS algorithm with values for these parameters or estimate them directly from the data. To estimate the transition probabilities from the data, we use the statistical definition of a probability. We also rely on pairwise alignments between biological isomiR sequences, *s*^′^, and isomiR sequences, *s*. Recall that here, pairwise alignment between biological isomiR sequence *s*^′^ and sequence *s* means the results of using the Needleman-Wunsch algorithm to obtain a global alignment of two nucleotide sequences (30).

Each type of transition from one specific character (nucleotide or gap) to another (nucleotide or gap if the starting character is a nucleotide; nucleotide if the starting character is a gap) defines an event of interest. If *s*^′^ and *s* are the biological isomiR and error isomiR sequence, respectively, and *l* indexes specific positions along the pairwise alignment of *s*^′^ and *s*, then an event *E* is described by the transition 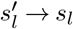. The sample space for the event *E* is the set of all possible transitions that can occur from the character 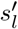. Then, the probability of observing event *E* is given by

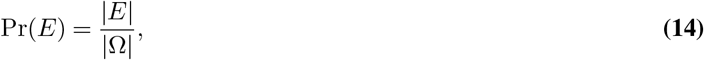

where |*E*| and |Ω| are the sizes of the sets representing the event *E* and the sample space Ω. To calculate the size of |*E*|, we must count the number of times the character 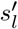 transitions to character *s*_*l*_ in any pairwise alignment between a biological isomiR sequence and an error isomiR sequence, for any miRNA in the profiling experiment.

## Supplementary Note 2: Simulating data from the error model

The proposed model for the process of generating error isomiR sequence reads from biological isomiR sequences gives us an assumed data-generating mechanism we can use to simulate data. The simulated data will consist of error isomiR sequences, generated from different biological isomiR sequences, mapping to different miRNAs and associated read counts. For each sequence *s* in the dataset, we will observe whether the sequence is a biological isomiR sequence or an error isomiR sequence. To generate simulated datasets that resemble a real, experimental dataset as much as possible, we need to generate isomiR sequences of different types i.e. length variant isomiRs, sequence variant isomiRs, mixed type isomiRs, etc. A single observation in our simulated dataset consists of the following: an isomiR sequence, *s*, mapping to miRNA *i*, with the mapping described by some function, *f, f*(*s*) = *i*, a read count *y*_*sj*_, and a label for the source of the sequence *s* as a biological isomiR sequence or a technical error isomiR sequence. We will describe the approach here in terms of generating observations for a single miRNA *i*. We begin with an experimental dataset so that we have real miRNAs. Furthermore, we use the experimental dataset for identifying biological isomiR sequences and associated read counts. This helps us to simulate a dataset that is as close to a real dataset as possible.

### Simulating length variant isomiRs

First, we estimate the transition probabilities in Θ according to the processes described in the previous section. Next, we select a miRNA *i* from the set of miRNAs present in the experimental benchmark dataset. We group all sequences *s* mapping to miRNA *i* in the set 𝒮_*i*_ = {*s* : *f*(*s*) = *i*} . Next, we identify the most abundant sequence *s* ∈ 𝒮_*i*_ as a true biological isomiR sequence for miRNA *i*. We will use 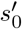 to refer to this biological isomiR sequence. Together, 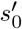 and its associated read count in sample *j*, 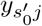 form the first observation in our simulated dataset.

The rest of the sequences *s* in our simulated dataset, mapping to miRNA *i*, will be generated by manipulating the text string representing 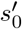 at random. To generate a length variant isomiR sequence 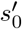, first we sample the difference in length between *s*^′^ 0 and the isomiR sequence *s* to be generated from the set {−7, −6, −5, −4, −3, −2, −1, −1, −2, −3, −4, −5, −6, −7}, with replacement. The definition of the Levenshtein distance between two strings is the number of single-character edits required to turn one string into the other. Therefore, if we are limiting ourselves to only adding or removing a single nucelotide, we can think of the integer we have sampled as a sampled value for the Levenshtein distance between 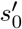 and *s*. We do not include 0 in the set from which to sample because it would result in the return of a sequence with no changes. We assume that each sequence appearing in the dataset is unique. We select 7 as the maximum number of differences we allow between 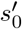 and *s* to prevent us from generating isomiR sequences that are too long or short to be identified as miRNAs by alignment software. Having isomiR sequences that are much longer or shorter than what would be identified in a real dataset moves the simulated data away from the ground truth, which is undesirable.

While the value we have sampled is similar to the Levenshtein distance, it is not the same. The Levenshtein distance is always an element of ℤ_*≥*0_. We allow negative integers in the set from which to sample because we the sign of the sampled value tells the user how the length of the simulated sequence differs from 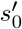. If the sampled value is negative, then error isomiR sequence *s* is shorter in length than 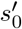. Otherwise, if the sampled value is positive, then error isomiR sequence *s* is greater in length than 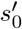. Let *L*_*d*_ be the sampled difference in length between 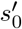 and the error isomiR sequence to be generated, *s*. Suppose *L*_*d*_ *<* 0. After sampling *L*_*d*_, we iterate through the set of integers from 1 to |*L*_*d*_|, where |*L*_*d*_| is the absolute value of *L*_*d*_. For each iteration, we sample an end of the isomiR sequence from the set {3*p*, 5*p*}, with repalcement. The elements of the set correspond to the 3’ end and 5’ ends, respectively of a genetic sequence. Let *e* represent the end we have sampled. Then, after sampling *e*, we remove a single nucleotide from the string representing 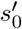 at end *e*. We continue this process for each iteration of sampling an end and removing the terminal nucleotide at that end until we have removed a total number of |*L*_*d*_| nucletoides from *s*^′^ 0. The resulting sequence *s* is a length variant isomiR sequence of 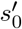 that is shorter than 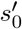. We label this generated sequence a technical error isomiR sequence.

Now, suppose *L*_*d*_ *>* 0. Again, we iterate through the set of integers ranging from 1 to |*L*_*d*_| = *L*_*d*_. At each iteration, we again sample an end *e* from the set *{*3*p*, 5*p}*, with replacement. However, because *L*_*d*_ *>* 0 is taken to represent a length variant isomiR sequence greater in length than 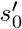, we must make additions to 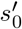 to generate *s*. Thus, in addition to choosing the end of the sequence to manipulate, we must choose the nucleotide to add from *{A, C, G, T }*. As we mentioned before, this corresponds to sampling adenine, cytosine, guanine, or thymine, respectively. We sample a nucleotide *z* from *{A, C, G, T }* with equal probability and replacement, to be added to end *e* of biological isomiR sequence 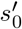. We continue this process of sampling an end *e* to make an addition to and sampling the nucleotide to add until we have added a total number of sampled nucleotides equal to *L*_*d*_. The result is an error length variant isomiR sequence *s* of 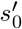 that is *L*_*d*_ nucleotides longer than 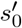.

Generating the sequences *s* and their labels are only part of the process. The next step to generate complete observations is to simulate read counts for the simulated sequences *s* from the error model. Consider a simulated isomiR sequence *s*, generated from sequence 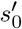, with read count 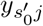 . First, we perform a pairwise alignment of *s* to 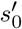. Next, we calculate 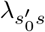 according to Equation 1. Then, we use 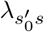 and 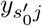 to calculate the error model distribution according to Equation 5. Finally, we draw *y*_*sj*_ from this error model.

### Simulating sequence variant isomiRs

Length variant isomiR sequences are but one type of isomiR sequence that exists. Another type of isomiR sequence we need to be able to simulate is sequence variant isomiRs. The process for making substitutions to a starting sequence 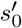 is similar to the process of making additions to a starting sequence 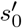.

Like with length variants, simulating sequence variant isomiRs begins with a real, experimental dataset. In the experimental dataset, we estimate the model parameters stored in Θ according to the procedure outlined in the previous section. For miRNA *i*, we form the set 𝒮_*i*_ = {*s* : *f*(*s*) = *i*}, that is all the sequences *s* mapping to miRNA *i*. Then, we select the most abundant sequence in 𝒮_*i*_ and call it 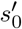. This acts as a single biological isomiR sequence for miRNA *i* in our simulated dataset. To generate a sequence variant isomiR sequence *s* from 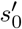, we begin by sampling the number of differences in sequence between 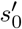 and *s* the isomiR sequence to be generated from {1, 2, 3, 4, 5, 6, 7}. Let *H* be the sampled difference in sequence between 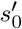 and *s*. The definition of the Hamming distance is the number of substitutions that need to be made to one string to generate another string of equal length. Here, because we specifically define sequence variant isomiRs as sequences of equal length to the reference sequence, *H* is equivalent to the Hamming distance between 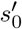 and *s*. We let *H* take on a maximum value of 7 to avoid generating error isomiR sequences that differ at too many points from the reference sequence. If such a sequence existed in the real data, it is highly probable that the alignment software would not align 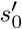 and *s* to the same part of the reference microRNAome.

To make a single substitution in 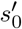, we must specify a position to make the substitution at and the nucleotide to use as the substitute. First, we sample the position *l* along the length of the sequence 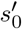. Let 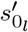 be the nucleotide at position *l* in 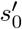. Then, after sampling *l*, we sample the substitute nucleotide from 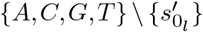 and make the change. We continue this process iteratively, first sampling a position, *l* along the length of the sequence 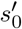 and then selecting a nucleotide to act as the substitute from the set 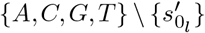 a total of *H* times. The resulting sequence *s* is a sequence variant isomiR of 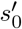 with a Hamming distance of *H*. We label *s* a technical error isomiR sequence. After generating the sequence variant isomiRs themselves, the process for simulating read counts of simulated sequence variant isomiRs is the same as the one used for simulated length variant isomiRs.

### Generating simulated datasets with multiple true biological isomiRs per miRNA

In our description of the methods for simulating length variant and sequence variant isomiRs above, we have begun by selecting a miRNA *i* from a real experimental dataset and identifying the most abundant sequence mapping to that miRNA. The result is a simulated dataset that has one biological isomiR sequence per miRNA. While useful from a statistical standpoint, this generates simulated datasets with characteristics that diverge slightly from experimental datasets. We know that there are multiple isomiRs per miRNA in an experimental dataset (34). Therefore, to generate simulated data with multiple biological isomiRs per miRNA, we first choose a miRNA *i* from an experimental dataset and then *N*_bio_, the number of biological isomiRs per miRNA we desire in the simualted dataset. Then, from the set 𝒮_*i*_ = {*s* : *f*(*s*) = *i*}, we select the *N*_bio_ most abundant sequences to act as the *N*_bio_ biological isomiRs in the simulated dataset. Then, we treat each of the *N*_bio_ biological isomiRs independently and follow the procedures outlined above to generate error isomiR sequences from a single biological isomiR sequence and associated read counts.

## Supplementary Note 3: Using *β* to create consensus sets of center sequences across samples

For miRNA *i*, PARiS infers partitions 𝒫_*j*_ of isomiR sequences mapping to miRNA *i* in each sample *j* = 1, …, *J*, independently. Each partition 𝒫_*j*_ is defined by a set of disjoint subsets, 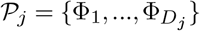, of sequences *s* mapping to miRNA *i* in sample *j*. Each subset Φ_*d*_ consists of an inferred center sequence, *ϕ*_*d*_, one of the inferred biological isomiR sequences mapping to miRNA *i* in sample *j*. The other elements of Φ_*d*_ are inferred to be error isomiR sequences of *ϕ*_*d*_, according to the error model. Therefore, the inferred composition of sequences for miRNA *i* in sample *j* is the set of center sequences defining the subsets of the partition 𝒫_*j*_. We use 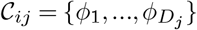 to denote the set of sample-specific center sequences for miRNA *i* in sample *j*. To infer the overall composition of sequences for miRNA *i* in the experiment, we need to combine the sets 𝒞_*i*1_,, 𝒞_*ij*_ into a single consensus set for miRNA *i*, 𝒞_*i*_.

In this manuscript we introduce a value *β*, used to generate the consensus set of center sequences 𝒞_*i*_. The value *β* controls the amount of pooling between samples, an alternative to taking the union of the sample-level sets of center sequences (no pooling) or taking the intersection of the sample-level sets of center sequences (complete pooling). *β* is a real-valued parameter that takes on a value from the interval 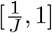, where *J* is the total number of samples in the miRNA profiling experiment. Setting 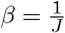 is the same as taking the union of the sample-level sets of center sequences and setting 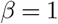 is the same as taking the intersection of the sample-level sets of center sequences.

For values of 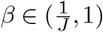, *β* is used as follows for creating a consensus set of center sequences: suppose for miRNA *i* we have sample-level sets of center sequences 𝒞_*i*1_, …, 𝒞_*iJ*_ . Let *s*^′^ be a center sequence from some 𝒞_*ij*_. To determine if *s*^′^ should join the set 𝒞_*i*_, we define a series of *j* = 1, …, *J* indicator variables as

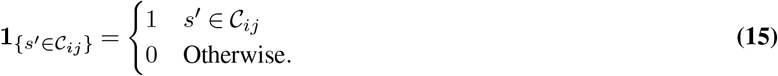

Then, for the center sequence *s*^′^, we calculate

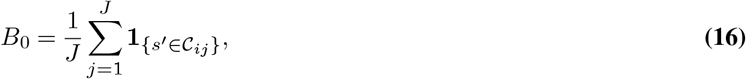

and compare it to *β*. If *B*_0_ *≥ β*, the center sequence *s*^′^ joins the consensus set of center sequences for miRNA *i*, 𝒞_*i*_. Otherwise, *s*^′^ does not join the set 𝒞_*i*_. After identifying all elements that should belong to 𝒞_*i*_, some subsets defining a partition may no no longer exists because the center sequence defining it is not in 𝒞_*i*_. For these sequences, we place it into the subset of sequences in that sample. After fully defining 𝒞_*i*_ and identifying all biological isomiR sequences *s*^′^ belonging to 𝒞_*i*_, and redefining set memberships as necessary, we have a partition of sequences mapping to miRNA *i* that can be applied uniformly to each sample *j* in the profiling experiment. Then, we can use the inferred sequence compositions for miRNA *i* as defined by the partition to generate a denoised version of the data for use in downstream analyses.

